# Filamentation and restoration of normal growth in E.coli using a combined CRISPRi sgRNA/antisense RNA approach

**DOI:** 10.1101/323212

**Authors:** Andrea Mückl, Matthaeus Schwarz-Schilling, Katrin Fischer, Friedrich C. Simmel

**Affiliations:** Physics of Synthetic Biological Systems (E14), Physics Department, Technische Universität München, Am Coulombwall 4a, 85748 Garching, Germany; Nanosystems Initiative Munich, 80539 Munich, Germany.

## Abstract

CRISPR interference (CRISPRi) using dCas9/sgRNA is a powerful tool for the exploration and manipulation of gene functions. Here we quantify the reversible switching of crucial cellular processes by CRISPRi and an antisense RNA mechanism. Reversible induction of filamentous growth in *E. coli* has been recently demonstrated by controlling the expression levels of the bacterial cell division proteins FtsZ/FtsA via CRISPRi. If FtsZ falls below a critical level, cells cannot divide. However, the cells remain metabolically active and continue with DNA replication. We surmised that this makes them amenable to an inducible antisense RNA strategy to counteract FtsZ inhibition. We show that both static and inducible thresholds can adjust the characteristics of the switching process. Combining bulk data with single cell measurements, we clarify the role of bacterial heterogeneity and population dynamics for gene circuits affecting cell division. Filamentation is shown to strongly increase gene expression variability in the bacteria. Furthermore, we find reversible switching only in a small subpopulation of the bacteria, which takes over the population upon continued cell division. Successful restoration of division occurs faster in the presence of antisense sgRNAs than upon simple termination of CRISPRi induction.

## Introduction

Bacterial growth rate and protein expression levels are tightly linked to the physiology of the cell (1). The control of physiological processes such as cell division or genome replication with artificial gene circuits therefore is of great interest for various applications in synthetic biology. For instance, growth control circuits could be used to switch between different growth modes to optimize metabolic processes.

As a recent example, Wiktor *et al.* controlled the growth of *E. coli* bacteria by preventing replication initiation of the protein DnaA using the CRISPR/dCas9 interference mechanism (CRISPRi) (2). To this end, the authors blocked the DnaA boxes at the *oriC* using sgRNAs of the appropriate sequence and the non-cleaving mutant of the CRISPR-associated protein 9 (dCas9) (3).

A different approach is to alter the growth characteristics of *E. coli* bacteria by inhibiting the expression of the bacterial tubulin homolog FtsZ, which is the key player of the bacterial divisome (4). It has been shown that the ratio of FtsZ to FtsA is a crucial parameter for the initiation of fission ring formation (5). An imbalance of FtsZ and FtsA results in cell division arrest and bacterial filaments that contain multiple copies of the bacterial chromosome (6). Recently, Elhadi *et al. (7)* and others (8) (9) have shown that CRISPRi can be used to reduce and restore FtsZ and MreB levels and thus change the morphology of *E. coli* cells. Filamentation also occurs naturally under a wide variety of conditions and is surmised to have important biological functions (10, 11). In this context, Sánchez-Gorostiaga *et al.* recently determined the physiology and recovery from FtsZ depletion with an IPTG inducible strain (12).

In the present work, we reversibly induce filamentation in *E. coli* by targeting FtsZ using the CRISPR/dCas9 interference mechanism combined with an antisense RNA strategy. We first switch bacteria into the filamentous cell growth mode and subsequently reverse this process, such that the cells return into a normal growth phenotype. We quantify the switching process and dynamically control the system by the inducible expression of antisense RNAs that are complementary to the sgRNAs engaged in *ftsZ* knockdown in comparison to the removal of the CRISPRi inducers alone. We use single-cell fluorescence microscopy experiments from which we derive bacterial length distributions (13), growth and division rates, as well as reporter gene expression levels. This allows us to identify factors that affect the switching process. In particular, we find that a subpopulation with low dCas9/sgRNA expression levels can be actively switched back to normal division by the induction of antisense RNAs (‘anti-sgRNAs’). The anti-sgRNA supported recovery process is found to occur considerably faster than switching in the absence of the antisense RNAs.

In order to reversibly switch bacteria to filamentous growth, we first disturbed the FtsZ/FtsA ratio through CRISPR interference using appropriate sgRNAs (Fig 1A). We then restored cell division using different strategies, including the use of antisense sgRNAs (Fig 1B).

Three of at least seven known promoters for *ftsZ* (ftsZ2p, ftsZ3p, ftsZ4p) lie within the *ftsA* coding region (14). We targeted these three promoters for transcription blockage using sgRNAs with the corresponding sequences (cf. Fig 1 and S1 Table). The sgRNAs are induced by IPTG via T7 RNA polymerase and were encoded on a plasmid together with TetR-controlled dCas9 and mVenus reporter protein (the ‘CRISPRi plasmid’).

In order to inhibit *ftsZ* repression inducibly, we use an anti-sgRNA plasmid, from which sequences complementary to the three sgRNAs could be transcribed. The anti-sgRNAs were designed to absorb the sgRNAs via duplex formation in a similar manner as previously demonstrated by Lee *et al.* (15). In our approach, the anti-sgRNAs are induced by AHL (the acyl homoserine lactone (HSL) 3-oxo-C6-HSL) from pLux promoters (cf. S2 Table). Both plasmids contain sgRNA ‘sponge’ elements (*vide infra*), which are decoy-binding sites for dCas9-sgRNA.

**Figure 1.**
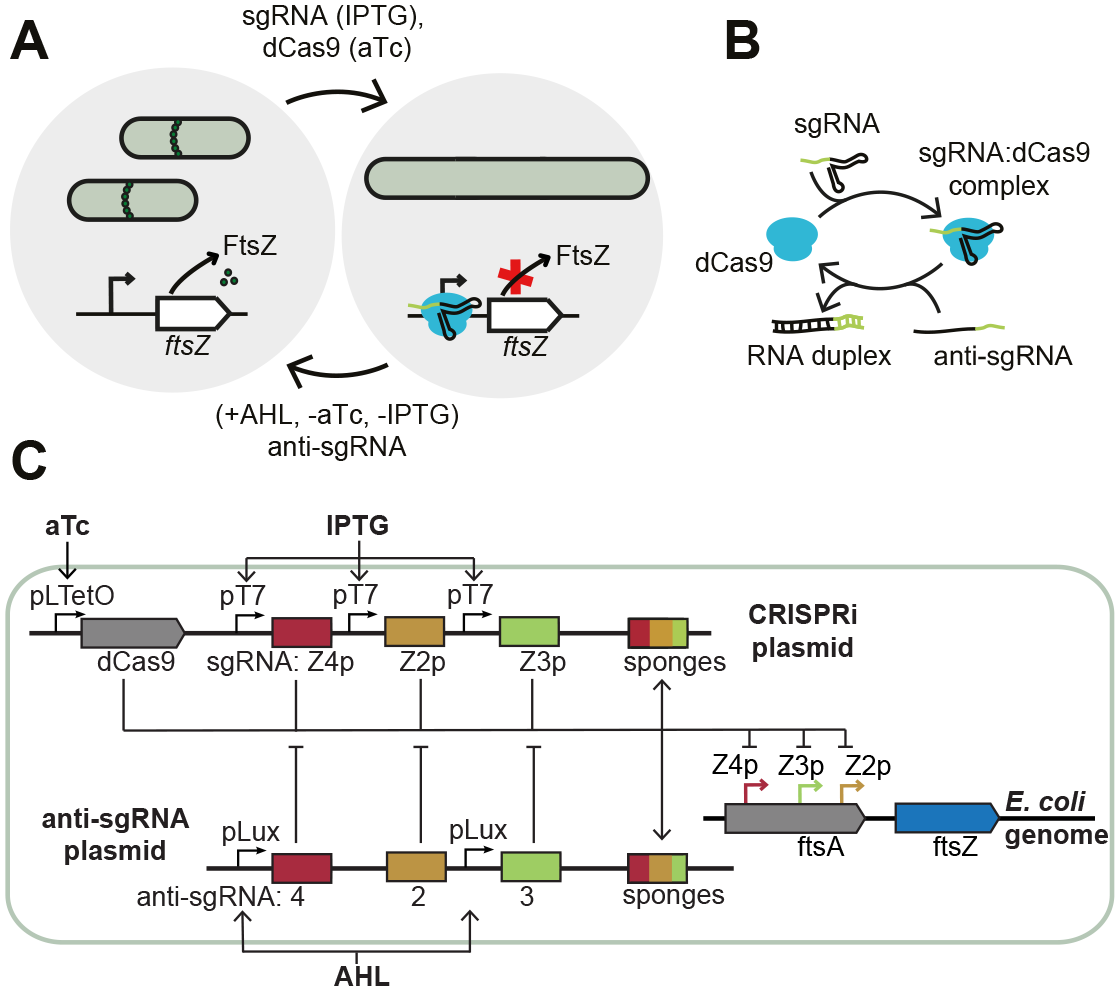
CRISPR-based growth control. **(A)**Schematic representation of *E. coli* switching into filamentous cells by inducing single guide RNA (sgRNA) via IPTG and dCas9 via aTc. The sgRNA-dCas9 complex blocks the expression of FtsZ, stopping the formation of the septal ring that is essential for cell division in *E. coli.* Cell division can be rescued by inducing appropriate antisense sgRNAs (‘anti-sgRNA’) with AHL and by removing the inducers for the dCas9 and sgRNA. **(B)** Scheme of sgRNA forming a complex with dCas9. The anti-sgRNA can inhibit the complex formation by binding to the sgRNA. **(C)** The involved genetic constructs in greater detail: the CRISPRi plasmid codes for dCas9 under aTc-inducible promoters and three different sgRNAs under T7 promoters which target three different promoters of the *ftsZ* gene on the genome of the *E. coli.* T7 RNA polymerase is inducible with IPTG. The ‘anti-sgRNA plasmid’ codes for anti-sgRNAs under the control of an AHL-inducible promoter. The sponge elements on the plasmids act as decoy binding sites for the corresponding dCas9-sgRNA complexes.

## Results and Discussion

### Decoy-binding sites to buffer CRISPRi

In contrast to tunable CRISPRi (tCRISPRi) with an inducible chromosome-integrated dCas9 (9), we here used plasmid-encoded dCas9 and decoy-binding sites (‘sponges’) to set the threshold for dCas9-sgRNA above which *ftsZ* is regulated down.

Thresholding was necessary to buffer away dCas9 in the presence of leaky expression of sgRNA from the T7 promoter. Transformation of bacteria with the CRISPRi plasmid alone resulted in filamentous cells (cf. Fig 2A). We therefore further increased the threshold for CRISPRi knockdown of *ftsZ* by introducing additional sponge elements also on the anti-sgRNA plasmid, upon which bacteria containing both plasmids exhibited the intended normal growth morphology (cf. Fig 2B & S1 Movie). Hence, sponge elements were required at a high copy number (anti-sgRNA plasmid ~500-700 per cell (16)), which is much higher than the copy number of the plasmid carrying the CRISPRi machinery (i.e., 20 ± 10 (17, 18)).

Expression of dCas9 in the absence of sgRNA or with sgRNAs of different sequence does not lead to filamentous growth under our experimental conditions. We can therefore rule out that dCas9 alone was responsible for the observed changes in cell morphology, as was reported previously for high level expression of the protein (19).

**Figure 2.**
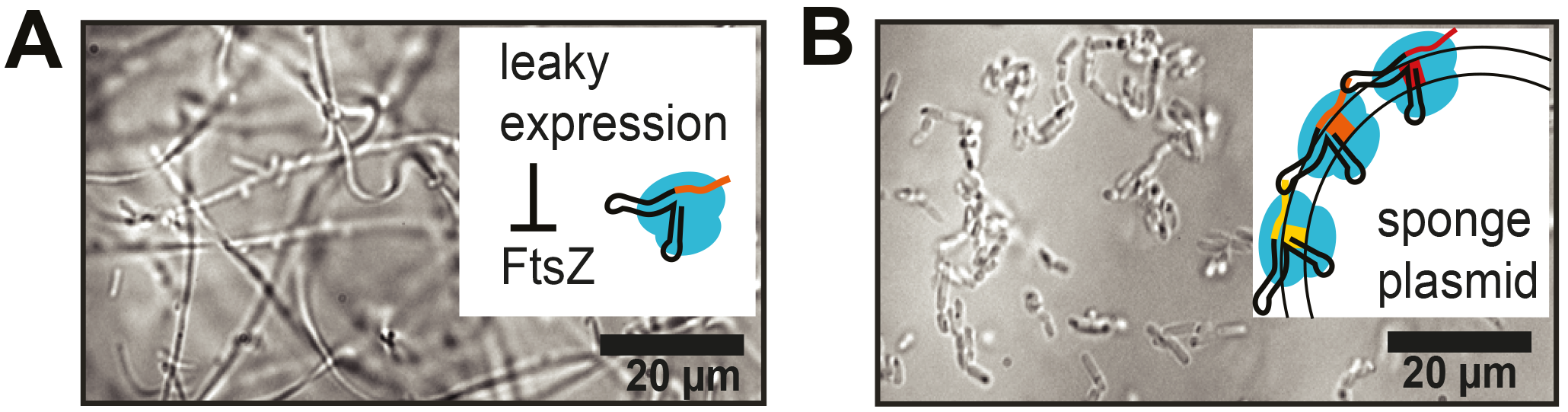
Decoy-binding site strategy. Bright field images of bacteria in liquid culture containing the CRISPRi-plasmid in the non-induced state without **(A)** and with **(B)** a high copy number of sponge elements.

### Reversal of CRISPR interference using antisense sgRNA

In order to test the effect of anti-sgRNA on CRISPR interference, we initially performed a series of *in vitro* experiments in bacterial cell extract (20). Here we regulated the expression of mVenus using purified dCas9 together with sgRNA and anti-sgRNA. As shown in Fig 3B, mVenus expression is efficiently suppressed in the presence of dCas9 and sgRNA (S3 Table for the corresponding sequences), while the addition of anti-sgRNAs recovers the production of mVenus. We tested two different designs for the anti-sgRNAs, which were complementary to part of the sgRNAs. Our experiments confirmed, that sequestration of the spacer region and only part of the dCas9 handle by anti-sgRNAs is already sufficient to de-activate the CRISPRi mechanism (Fig 3A-C). In the *in vivo* experiments, both anti-sgRNA designs were used together.

In a further control experiment, we checked the FtsZ levels by immunoblotting. CRISPRi efficiently decreases FtsZ expression in *E. coli*, while anti-sgRNA induction recovers the FtsZ concentration despite the presence of inducers (Fig 3D).

**Figure 3.**
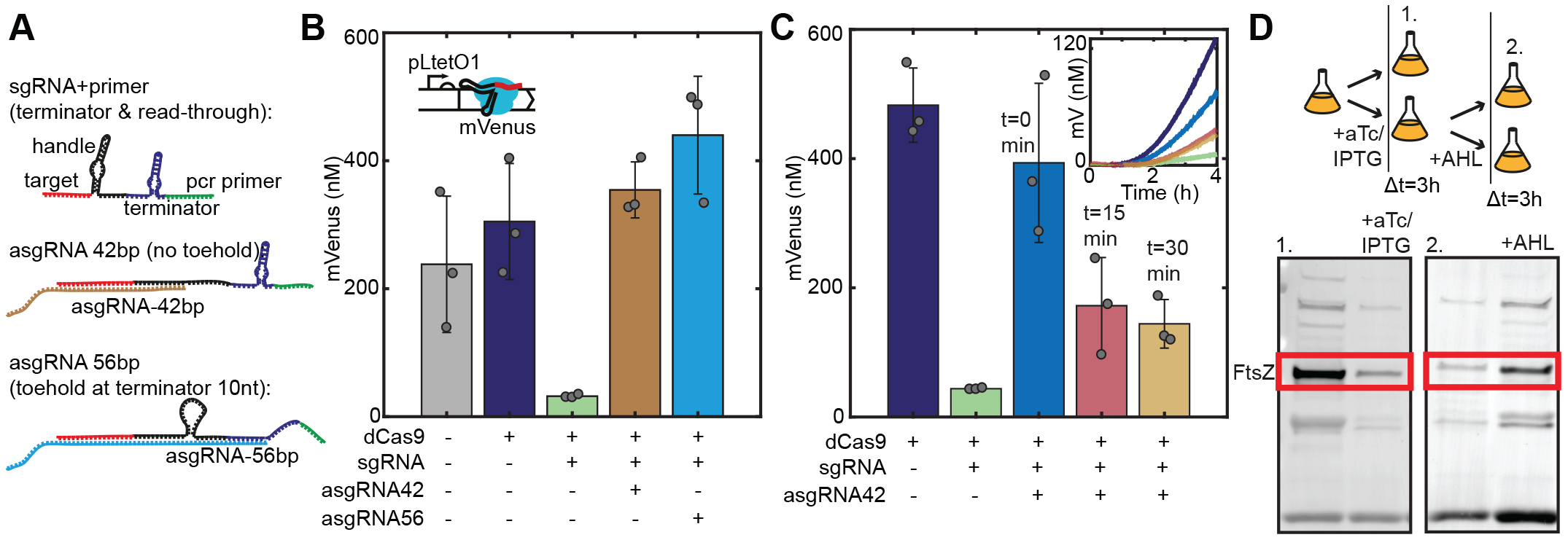
Anti-sgRNA strategy tested *in vitro*. **(A)** Secondary structures of free and complexed sgRNA:asgRNA variations. The dCas9 handle is destroyed by duplex formation, which prevents dCas9 from binding **(B)** Prototyping the CRISPRi knockdown and the rescue system in a cell-free transcription/translation system. The restoration of mVenus signal is shown with truncated anti-sgRNA versions (the number indicates the number of base pairs in the resulting sgRNA:asgRNA duplex). The expression of mVenus ([template DNA] = 5 nM) is blocked by the supplementation of purified dCas9 (70 nM) and sgRNA (100 nM) and is re-activated upon addition of anti-sgRNA (0.5 μM). Fluorescence levels for three different samples are taken at t = 15.5 hours or t = 11 h. **(C)** Delaying the time of anti-sgRNA addition relative to dCas9-sgRNA results in lower mVenus fluorescence intensities. The inset shows the fluorescence time traces corresponding to the fluorescence intensity end levels shown in the main panel (taken at t = 12 hours). Error bars are plotted as SD from 3 replicates (c(sgRNA) = 250 nM, c(asgRNA) = 1μM). **(D)** Immunoblot showing FtsZ levels of normal and filamentous (+46 nM aTc, +215 μM IPTG) cells. Diluting and splitting the filamentous cell culture with and w/o AHL (in the presence of aTc and IPTG), shows the recovery of FtsZ in the presence of AHL (50 nM).

### Single-cell analysis of filamentation

After induction of the CRISPRi mechanism with aTc and IPTG, bacterial cells rapidly stopped to divide and started the expression of mVenus (which also was under the control of a pLTetO promoter). We performed time-lapse video microscopy studies (cf. Materials and Methods) to observe over time the growth and fluorescence of individual filamentous bacteria inside microfluidic trap chambers that are connected to fresh medium supply channels (S1 Fig). The cell length distribution of *ftsZ*-knockdown bacteria broadens and shifts towards greater lengths (Fig 4A). The mean cell length increases from <L>=3 μm to <L>=21 μm in three hours with 500 μM IPTG and 107 nM aTc (which we defined as the 100 % induction level). As found by Li *et al.* (9), at low induction levels there is a co-existence of subpopulations of normal growing cells and filamentous ones. Upon 100 % induction of the CRISPRi mechanism, we were able to switch to filamentous cell growth (with a division rate approaching zero, cf. inset of Fig 4A). As shown in Fig 4B the average length of the bacteria increased faster with higher inducer concentrations. The filamentous growth of *E. coli* proceeded for up to 10 hours after which all the bacteria finally burst (S2 Movie).

**Figure 4.**
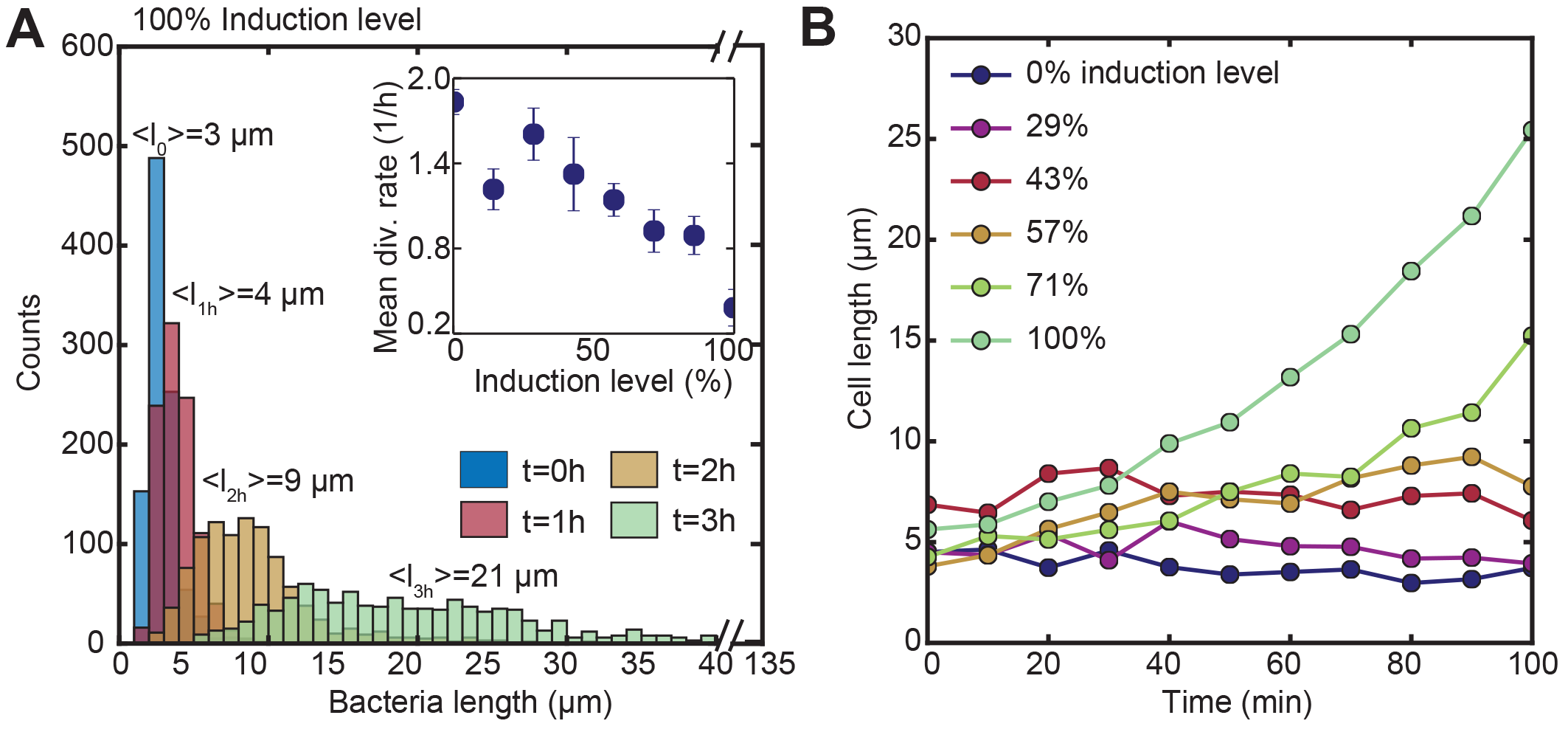
Filamentous bacteria. **(A)** Histogram of the cell length (calculated from 1000 cells each) at different time points at 100 % induction level (corresponding to 500 μM IPTG and 107 nM aTc). Over time the distribution broadens and the whole population shifts to larger lengths. Division rates decrease with increasing inducer concentrations. Results are means over 100 min. (for inset figure). **(B)** Tracking the cell length at different induction levels revealed a faster increase of the average cell length for higher induction levels.

### Efficiency of restoration of normal cell division

We investigated three different strategies to revert the filamentous cells to normal growth: i) either passively by stopping the production of dCas9/sgRNAs via removal of the inducers aTc and IPTG, ii) by induction of the generation of anti-sgRNAs actively counteracting the CRISPRi mechanism (the ‘antisense’ strategy), or iii) by a combination of both methods (the ‘active’ approach) (Fig 5A).

Interestingly, the three strategies performed differently in different environments. In bulk experiments with bacterial cultures in flasks, all three strategies allowed us to switch the bacteria back to normal growth.

As shown in Fig 5B for the passive and the antisense strategy, the cell length distribution shifted from about <L> = 10 μm to <L> = 4 μm within a few hours. A flow cytometer experiment performed for the active strategy showed that upon IPTG/aTc induction about 93 % of the cells switched to higher mVenus fluorescence intensity and higher forward scatter (FSC) signal (Fig 5C & S2 Fig). After supplementation of AHL (60nM for 3 hours) the non-filamentous population increased again from 6 % to 28 %.

The used AHL concentration did not affect growth as shown in a control experiment (S3 Fig).

In contrast to the bulk experiments cells that were induced to grow filamentous inside microfluidic chambers usually did not re-initiate cell division under strategy ii), i.e., upon supplementation of AHL and in the presence of aTc and IPTG (S4 Fig) (the only exception, where strategy ii) was successful in microfluidics, is shown in S3 Movie). Only the removal of the CRISPRi inducers allowed a fraction of about 5 % of > 2500 analyzed cells to resume normal cell division (S4 Movie).

Strategy ii) requires the sequestration of sgRNAs by anti-sgRNAs in the presence of dCas9 – thus dCas9 and anti-sgRNA will compete for binding to the sgRNAs present. The antisense strategy therefore will work better for lower dCas9 concentration and, accordingly, higher anti-sgRNA concentration. As these components are expressed from two different plasmids with different copy numbers, there may be large variations in the anti-sgRNA/dCas9-sgRNA ratio within a population (cf. also next section). The difference between the bulk and microfluidics experiments therefore can be interpreted as the result of simple statistics.

In all three strategies, restoration of normal growth occurred within a few hours, which is however slower than in studies, where *ftsZ* was directly put under the control of an inducible promoter (12).

**Figure 5.**
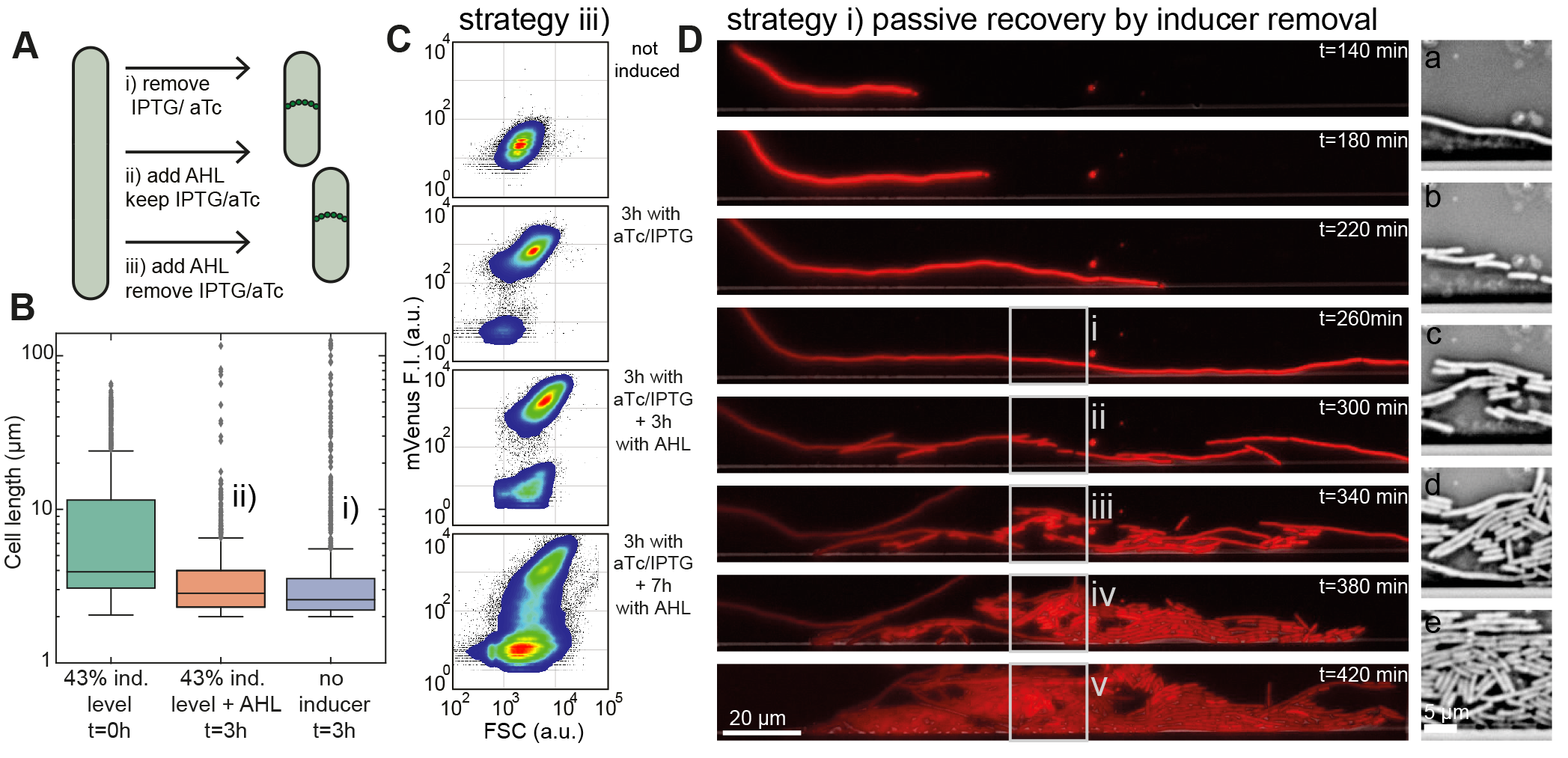
Restoration of cell division. **(A)** Scheme showing three different ways by which filamentous bacteria can be returned to normal cell division and growth. **(B)** Box plot of cell lengths (1000 cells each) of induced filamentous cells (at t = 0 hours) and their length after 3h in medium supplemented with AHL (in the presence of IPTG, aTc), or after 3 hours in medium without any inducers. For both rescue strategies, the mean cell size shifted back from <L> = 10 μm to <L> = 4 μm. **(C)** Flow cytometer density plots of bacteria displaying mVenus fluorescence vs. forward scatter (FSC) signal from a non-induced, normal growth state (top) towards the filamentous state by induction with IPTG/aTc (second) and back again (bottom) by supplementation of AHL (in the presence of IPTG/aTc). The mVenus and FSC signal increase in the filamentous growth mode. After the release of anti-sgRNA, the filamentous population decreases and turns back to normal cell size characteristics. **(D)** Left column: Fluorescence microscopy time series in the mRFP channel of an *E. coli* cell in a microfluidic chamber in the absence of inducers. Prior to the first image (top), the bacterium has been grown filamentous for two hours at 43 % induction level followed by 140 minutes exposure to growth medium without any inducers. At 300 minutes of growth, the bacterium resumes cell division. The new daughter cells quickly approach the normal cell size. Right column: images (a)-(e) show bright field images corresponding to the boxed regions on the left.

### Transient phenotypic heterogeneity

Our experiments in microfluidic cell traps revealed a considerable phenotypic heterogeneity upon induction of filamentation, which is reversed once normal growth is restored. In addition to their size, we characterized the cells with respect to their fluorescence intensity (Fig 6A & B). After 2 hours of continuous induction with IPTG/aTc, the coefficient of variation (CV=standard deviation/mean) of the cell size as well as of mVenus and mRFP expression increased considerably (Fig 6A). In contrast to the aTc-induced mVenus (CRISPRi plasmid), the constitutively expressed mRFP (anti-sgRNA plasmid) level dropped during filamentation (Fig S5). The decrease in mRFP fluorescence intensity could be caused by a dilution of the plasmid, a reduced protein expression rate or both. The same holds for the increasing CVs, which presumably are caused by greater variations in the plasmid copy numbers in the larger cells and/or in protein expression strength. Although both plasmid copy number controls are based on a negative feedback mechanism that measures the concentration of the plasmids in the cells (21), it has been shown that the copy number can change considerably depending on the bacterial growth rate (22). After removal of the inducers, a new population of normal growing cells restored similar CV values as the starting population.

**Figure 6.**
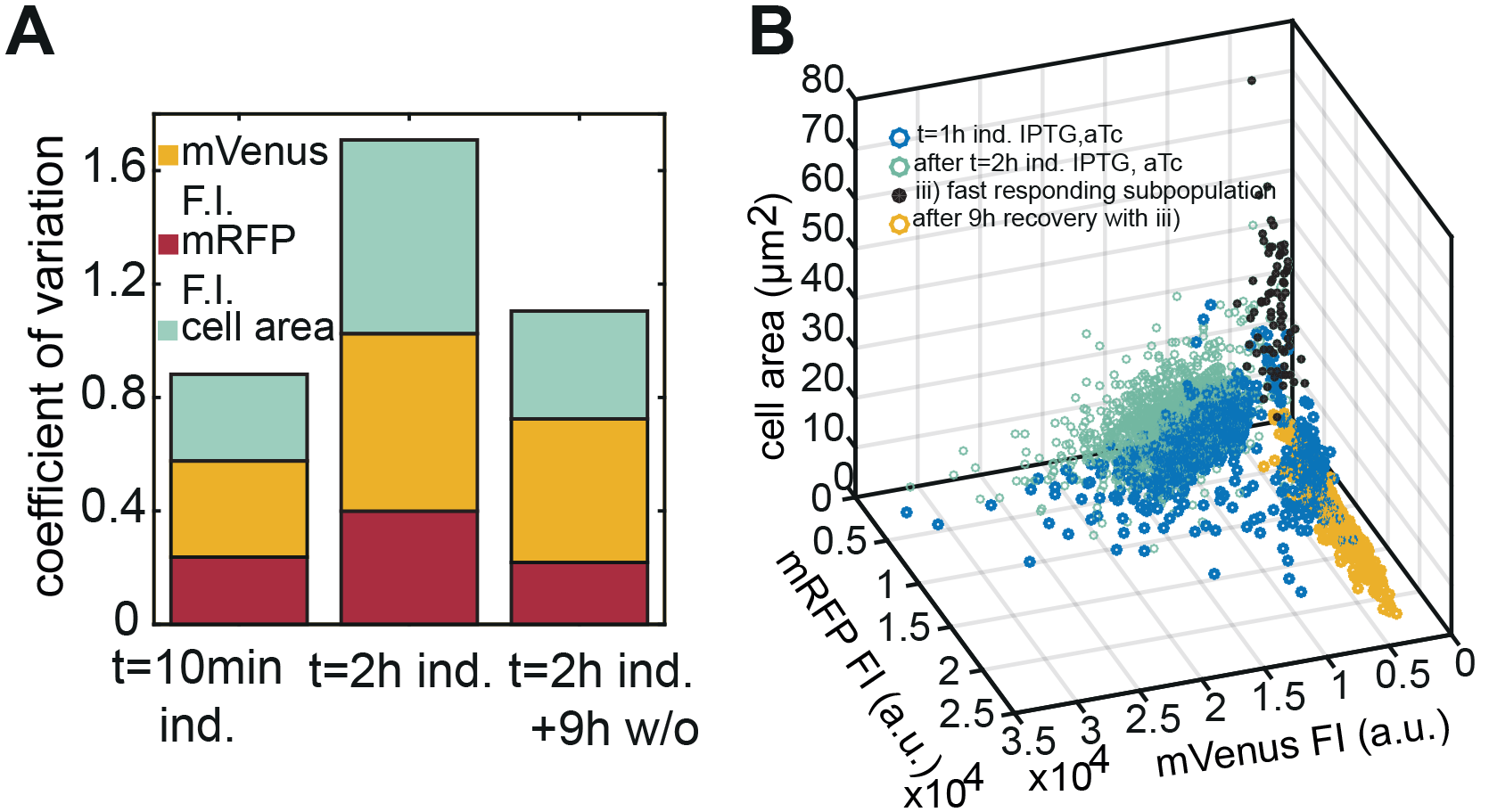
Single cell analysis of cell division switching. **(A)** The coefficients of variation (C.V.) were calculated for the fluorescence intensities of mVenus and mRFP and the cell area. The variability of the population increases during the 2 hours induction period and drops again for the new population emerging by passive switching, i.e., after removal of IPTG/aTc. **(B)** 3D scatter plot of an actively switched bacterial population (switching supported by anti-sgRNA in strategy iii). About 5-6 % of the population reinitiates division upon removal of IPTG/aTc from the growth medium while supplementing AHL (60 nM) in order to induce the expression of anti-sgRNA (‘fast dividers’: black dots).

### Inducible restoration of normal cell growth in a subpopulation

In contrast to the passive switching strategy described above (Fig 5D & S5), active switching according to strategy iii) switches back a different subpopulation of the filamentous bacteria (Fig 6B & S6). This subpopulation is characterized by low mVenus and mRFP fluorescence levels (Fig 7A & B) and a large cell size (Fig 6B). The mean mVenus fluorescence of a fast responding bacteria is 0.2 times lower than the average of the main population.

Low mVenus fluorescence should correlate with low dCas9 concentration (mVenus and dCas9 on the CRISPRi plasmid are both under TetR-control), which suggests that only in this case the expression of anti-sgRNAs can restore FtsZ levels enough to restart cell division.

Re-initiation of AHL supported cell division on average started about 80 minutes after removal of IPTG/aTc, which is ≈ 180 minutes faster than in the absence of AHL (Fig 5D & 7C). This means that in the confined space of a microfluidic growth chamber actively switched cells can take over the population before the other subpopulation (only responding to IPTG/aTC removal) switches back. The median length of the cells at first division was about 10 μm long (S6 Fig).

**Figure 7.**
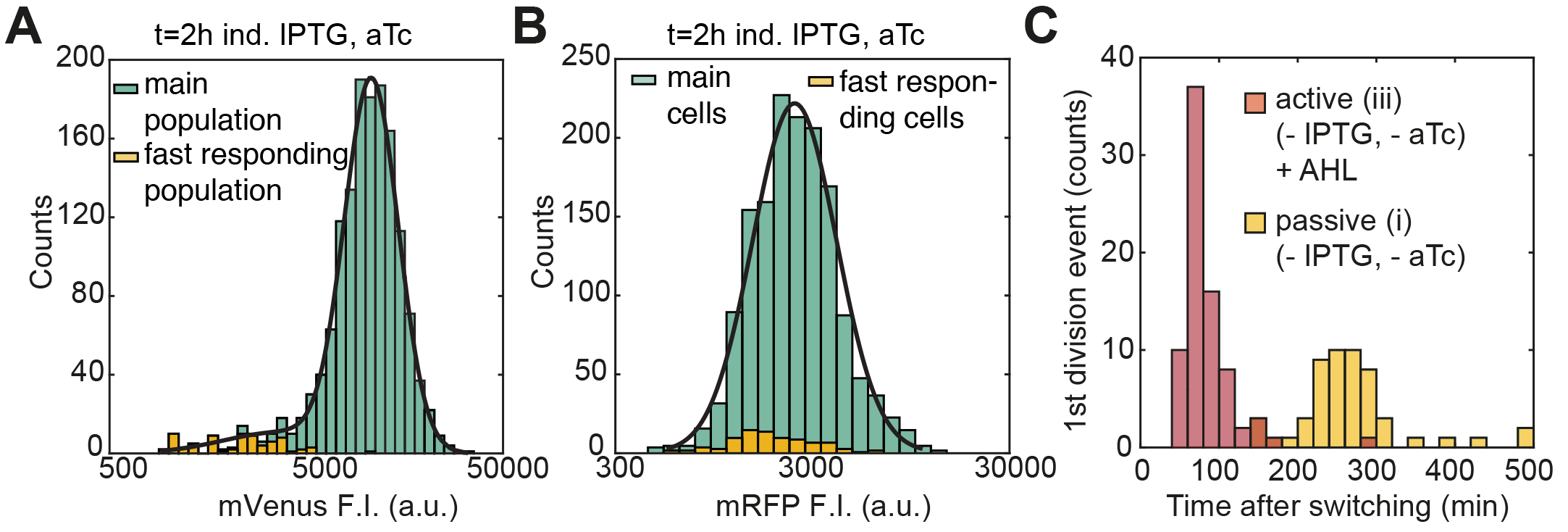
Characteristics of antisense RNA switchable cells. The fast switching subpopulation is characterized by low mVenus **(A)** and medium mRFP levels **(B)** (yellow bars) after two hours of induction of filamentous growth in microfluidics. The mVenus has been fitted with a Gaussian mixture model for two populations and mRFP with one population (fits are shown as solid black lines), **(C)** Histogram of the time passed between the removal of IPTG/aTc with and without AHL at the first cell division event (<t_+AHL_> ≈ 80 min, <t _-AHL_> ≈ 260 min). Actively reverted cells recover faster than passively switched cells.

## Conclusion

Using CRISPR interference and anti-sgRNA elements, we implemented a synthetic gene circuit that enabled us to controllably switch cell division off and on. Cells could thus be switched from normal cell division into a filamentous state, and also back to normal cell division.

Due to the critical role of cell division in the bacterial life cycle, our circuit strongly interferes with the physiology of the bacterial host chassis. In order to be able to implement the circuit in viable bacteria at all, decoy-binding sites for the FtsZ-suppressing dCas9-sgRNA complexes had to be introduced as genetic buffer elements. Next to these static sgRNA sponges, we also utilized inducible antisense-sgRNA molecules that could be used to adjust the threshold for bacterial filamentation and thus modulated the switching process.

An important aspect of a gene circuit that interferes with bacterial growth is that it very directly alters bacterial fitness. As a consequence, the circuit creates distinct dynamics both at the cellular and at the population level, which furthermore depends on the specific environment in which the bacteria grow. Using microfluidic single cell experiments, we observed that in this context only a fraction of the bacteria displays the desired switching behavior. However, these correctly switching bacteria can quickly take over the population, whereas others are expelled from the microfluidic chambers or die. Thus, on the population level the circuit appears to behave as designed.

In this context, it is interesting to note that upon induction of filamentation, the heterogeneity of our bacterial population appeared to increase considerably, and, depending on the switching strategy, different subpopulations of this population were selected. One interpretation of this observation is that the displayed heterogeneity is part of a bacterial strategy to survive the conditions imposed upon them by the synthetic circuit (23, 24).

Even though the response of the bacteria to a gene circuit directly affecting bacterial division or growth thus appears to be complex and quite problematic from an engineering perspective, achieving control over physiological processes is still highly desirable. The conflict of interest between the engineer’s agenda to produce a metabolite and the bacterial agenda to replicate (25), could be solved, in principle, by establishing molecular programs that switch between different cellular growth modes allowing an appropriate shift in resources (1, 26, 27). Such programs will inevitably have to take into account similar effects of bacterial heterogeneity and population dynamics as those described here.

## Materials and Methods

### Plasmids

We constructed plasmids by Gibson assembly of synthetic DNA fragments into the target vector (pSB1K3 and CRISPRi plasmid 44249 from addgene). The final plasmid sequences can be found in S2 Table. The sender strain was constructed in an earlier study and contains the gene for LuxI synthase (BioBrick part BBa_C0261). The sgRNA, anti-sgRNA and sponge element sequences are listed in S1 Table.

### Bacterial cell culture

Experiments with filamentous cells were performed in *Escherichia coli* BL21 (DE3) pLysS. Cells from glycerol stock were grown in 5 ml Luria-Bertani medium containing antibiotics selecting for both plasmids (CRISPRi and asgRNA plasmid) and incubated over night at 37 °C and 250 rpm. The following day, cells were diluted 1:100 and incubated for additional 2 h. Optical density (OD) values between 0.4-0.6 were obtained. From this batch, 1 ml of the culture was centrifuged and the pellet resuspended in 300 μl growth medium. The concentrated cells were immediately loaded on a microfluidic chamber until single or few bacteria were captured in the traps. In such a microfluidic chemostat with defined bacterial trap dimensions, we supplied the bacterial suspension constantly with fresh nutrients (LB medium, antibiotics and/or inducer chemicals or dyes) using a pressure flow controller (OB-1K, Elveflow).

### Fluorescence time-lapse microscopy

The microfluidic PDMS (Sylgard 182, Dow Corning) device was fabricated using standard soft lithography as previously described (28). The microfluidic device is a combination of a gradient mixer (29) and bacterial traps designed with dimensions of 200 μm × 10-50 μm × 1 μm as a H-shaped chemostat (S1 Fig).

Time-lapse microscopy measurements were conducted on a Nikon Ti-Eclipse epi-fluorescence microscope controlled with NIS-Elements Imaging Software. The microscope was equipped with a sCMOS camera (Zyla, Andor), an automated x-y-stage (Prior Scientific, Cambridge, UK) and an incubator box (Okolab) to maintain an operation temperature of 37° C. All videos were recorded with 40x apochromatic magnification objectives. Every 5 to 20 min, images in phase contrast mode, YFP as well as RFP fluorescence mode (in combination with the appropriate filter sets) were taken for a total run time of up to 20 hours. The exposure times were automatically adjusted.

### Cell-free expression

For the cell-free assay, the sgRNA sequences were designed complementary to the non-template strand (sequences can be found in Supporting Information S3 Table). Anti-sgRNA and sgRNA where transcribed *in vitro* by T7 RNA Polymerase (NEB) from linear DNA (IDT) overnight and then extracted with Phenol-Chloroform. The concentration of the RNA was determined by comparing a SYBR Green II stained band in a denaturing PAGE (8M Urea at 45°C) to the RNA Ladder (NEB, N0364S). The plasmid with mVenus was purified using Phenol-Chloroform prior the reaction in the cell extract. The crude S30 cell extract was obtained by beat beating of a BL21-Rosetta2(DE3) mid-log phase culture with 0.1 mm glass beads in a Minilys device (Peqlab) and supplemented with an energy mix and reaction buffer as described in ref. (20). Instead of 3-phosphoglyceric acid (3-PGA), phosphoenolpyruvate (PEP) was utilized as an energy source. dCas9 was His-tagged and purified by gravity-flow chromatography with Ni-NTA Agarose Beads (Qiagen). The fluorescence intensity was measured with a FLUOstar Omega plate reader (BMG) in 96-well plate (ibidi) at 37°C. The composition of a single cell-free reaction was: 33 % (v/v) S30 cell extract mixed with 42 % (v/v) buffer and 25 % (v/v) DNA plus inducers.

### Western blotting

An overnight culture was diluted in LB to OD 0.25 and grown for three hours with and without inducers (107 nM aTc and 1 mM IPTG). The sample with inducers was split and diluted again (to OD 0.25); one with 107 nM aTc and 1 mM IPTG and the other with 107 nM aTc, 1 mM IPTG and 100nM AHL. Before and after the dilution, 2 ml from each sample were pelleted and suspended in lysis buffer (50mM Tris, 14mM MgGlu, 60mM KGlu, 1mM DTT, 0.1 % TritonX100, pH 7.7) so that each sample had a concentration of 5 μg/ml (calculated from the OD). The samples were lysed with sonication on ice, pelleted and the supernatant was denatured at 95°C in Lämmli buffer and resolved by sodium dodecyl sulfate-polyacrylamide gel electrophoresis. 20 jg of proteins were transferred to PVDF membranes using a semi-dry transfer-blot apparatus (Bio-Rad). The membranes were blocked with 5 % (w/v) BSA in TBST overnight at 4°C and probed with anti-sera to FtsZ (1:1000, Agrisera) for 1,5 h at room temperature (30). A TRITC anti-mouse secondary antibody (1:5000; Agrisera) was applied for 1h at 4°C in 5 % BSA in TBST for 1h and the blot was imaged with a Typhoon FLA 9500 scanner (General Electric) in the Cy3 channel.

### Flow cytometry

An overnight culture was diluted (1/100) to 5 ml culture in LB and supplemented with appropriate antibiotics. The cell suspension was incubated for 2 hours and then induced with 100 μm IPTG and 11 nM aTc. 1 ml cell suspension was centrifuged and the pellet was solved in 2 ml PBS. For measurements with a flow cytometer (Cube8, Partec), the sample was further diluted with PBS (1:3). The anti-sgRNA was induced with 60 nM AHL for 3h and 7h before measuring. Cultures were diluted every 3 hours to keep the bacteria in the exponential growth phase. For each experiment, 100.000 – 150.000 events were recorded in FSC, SSC and FL1 (488ex/536em) and FL2 (532ex/590em) mode for mVenus and RFP detection.

### Data analysis

Image analysis was performed using NIS-Elements (Nikon) and customized MATLAB software. Flow cytometry data were plotted with FlowJo.

## ACKNOWLEDGMENTS

We gratefully acknowledge financial support by the BMBF through the ERASYNBIO network (project UNACS, grant no. 031L0011). M. S.-S. acknowledges support by the DFG Research Training Group 2062. We would like to thank Tobias Pirzer for useful discussions and Katharina HauUermann for experimental assistance with the Western blot.

## SUPPORTING INFORMATION

Additional movies, figures and tables based on descriptions in the text. Figures and tables are provided in a single “SI.pdf” file.

**S1 Movie.** This video shows *E. coli* (with the CRISPRi and anti-sgRNA plasmids) in a microfluidic chamber without inducers of the CRISPRI mechanism. The images are an overlay of BF/phase contrast and fluorescence channels of mVenus and mRFP. Time is shown as hh:mm.

**S2 Movie.** This video shows filamentous growth of *E. coli* in microfluidic chambers upon induction with 500 μm IPTG and 107 nM aTc (100 % level). The images are an overlay of BF/phase contrast and fluorescence channels of mVenus and mRFP. Time is shown as hh:mm.

**S3 Movie.** Active switching in microfluidic chambers. Filamentous growth is induced (215 μm IPTG and 46 nM aTc) for 2 hours. From there on the freshly supplied medium does not contain IPTG and aTc, but is supplemented with 50 nM AHL. The video starts after 1 hour of induction. One cell starts to re-divide about 50 minutes after the medium change. The images are an overlay of BF/phase contrast and fluorescence channels of mVenus and mRFP. The time is shown in hh:mm.

**S4 Movie.** Passive switching in microfluidic chambers. Filamentous growth is induced (215 μm IPTG and 46 nM aTc) for 2 hours. After the time window, the freshly supplied medium is without inducers. The images are an overlay of BF/phase contrast and fluorescence channels of mVenus and mRFP. This video shows a bacterium that has a relatively low growth rate during induction and takes relatively long to start re-division. The time is shown in hh:mm.

**S1 Figure. Microfluidic trap dimensions for single cell measurements.**

**(A)** The medium inlet direction is indicated with arrow marks. The exchange of nutrients and waste products occurs via diffusion. The channel width and trap dimensions are given in the layout. **B)** The cannel height is 15 μm whereas trap height is 1 μm. The traps are incorporated in a microfluidic gradient mixer (29).

**S2 Figure. Quantitation of the switching process using flow cytometry.**

The mRFP (top panel) and the SSC signal (bottom panel) are plotted against the FSC signal. mRFP is constitutively expressed in initially normal cells but also after induction of filamentation. After switching actively back with AHL in the presence of filamentation inducers IPTG/aTc for 3h, a population with decreased mRFP signal arises, which finally represents the main population (after 7h of asgRNA induction). In the filamentous state, the SSC signal ramps up in proportion with the FSC signal (t=3h with aTc/IPTG). However, rescued cells quickly recover and the majority of the cells turn back to the initial scatter plot position as measured with normal growing and dividing bacteria (cf. not induced and after t=7h with aTc/IPTG/AHL).

**S3 Figure. Cell growth and expression of fluorescent protein in bulk measurements.**

**(A)** Bacterial growth measured by monitoring the absorbance of the bacterial culture at λ=600 nm for various inducer concentrations. Bacterial growth was not affected by the induction of filamentation with aTc and IPTG nor by the supplementation of AHL. Three samples for each induction level are shown. **(B)** Fluorescence intensity time traces of mVenus measured at λem = 540 nm (excitation at 540 nm). **(C)** Fluorescence intensity of mRFP measured at λem = 590 nm (excitation at 544 nm). The fluorescence signal obtained from mRFP was both delayed and reduced for samples with AHL.

**S4 Figure. Cell division in the presence of inducers IPTG/aTc/AHL.**

The images are an overlay of bright field (phase contrast) and the fluorescence channels of mVenus and mRFP. During the first 1 hour and 20 minutes the cells are exposed to 215 μM IPTG and 46 nM aTc (43%). After 1 hour, 60 nM of AHL is supplemented to the growth medium. One bacterium resumes cell division (marked by an arrow), which belongs to a persisting subpopulation. The time is shown in hours. Scale bar: 20 μm.

**S5 Figure. Passive switching in microfluidic chambers.**

**(A)** 3D scatter plot with the mean mRFP and mVenus fluorescence levels of single bacteria plotted against the observed area of the corresponding cells for about 1000 cells at three different points in time: 10 minutes after induction (43% induction level), after 2 hours of continuous induction and after additional 9 hours without aTc/IPTG. About 5% of the filamentous cells divide (‘dividers’: black dots) again after the inducers have been removed from the growth medium. **(B)** Upon aTc/IPTG induction, the cells shift towards higher mean mVenus fluorescence intensities, lower mean mRFP and higher mean cell area. After 9 hours without inducers the mean values return to the level of the first measurement (t=10 min ind).

**S6 Figure. Active switching in microfluidic chambers.**

**(A)** Histogram of cell lengths. Actively switched cells (strategy iii) regain normal cell length distributions similar to t = 0 hours of starting induction of filamentous growth (cf. Fig 4 main text). **(B)** Histogram of the lengths of the daughter cells after the first division.

**S1 Table. Sequences of target sites, sponge elements, sgRNAs and anti-sgRNA.**

DNA sequences of the target sites of *ftsA* (Gene ID: 944778)) and the derived elements employed in this study.

**S2 Table. Plasmid sequences and description.**

The table shows the plasmid features of the constructed CRISPRi plasmid, the sponge plasmid and the anti-sgRNA plasmid in detail.

**S3 Table. Sequences for the cell-free assay.**

The DNA regions of interest in this study are summarized here.

